# Are female scientists underrepresented in self-correcting science for honest error?

**DOI:** 10.1101/2022.10.11.511375

**Authors:** MD Ribeiro, J Mena-Chalco, KA Rocha, M Pedrotti, P Menezes, SMR Vasconcelos

## Abstract

Retractions are among the effective measures to strengthen the self-correction of science and the quality of the literature. When it comes to self-retractions for honest errors, exposure of one’s own failures is not a trivial matter for researchers. However, self-correcting data, results and/or conclusions has increasingly been perceived as good research practice, although rewarding such practice challenges traditional models of research assessment. In this context, it is timely to investigate who have self-retracted for honest error in terms of country, field, and gender. We show results on these three factors, focusing on gender, as data is scarce on the representation of female scientists in efforts to set the research record straight. We collected 3,822 retraction records, including research articles, review papers, meta-analyses, letters, and others, from the Retraction Watch Database (2010-2021). We screened the dataset collected for research articles retracted for honest error, excluding retractions by publishers, editors, or third parties, and those mentioning any investigation issues. We then categorized the records according to country, field, and gender, after selecting research articles with a sole corresponding author. Our results show that female scientists account for 25% of self-retractions for honest error, with the highest share for women affiliated with US institutions.

## 1 Introduction

Retractions are among the effective measures to strengthen the self-correction of science and thus the reliability and quality of the literature. Concerning self-retractions for honest errors, whereas exposure of one’s own failures is not a trivial matter for researchers, self-correcting data, results, and/or conclusions for unintended errors has increasingly been perceived as good research practice (ALLEA, 2017; Global Research Council, 2021; Ribeiro et al., 2022). Recognition for such practice, however, is not (yet) part of the culture of rewards in academia (Nature Human Behavior, 2021; Bishop, 2018). One reason is that mechanisms to correct the literature, with post-publication explanations for invalidating, for example, data and/or conclusions of a research article, including self-retractions for honest error, gained traction only in the last two decades (Fang et al., 2012). Another reason is that those leading science today have built their careers within a culture of rewards based mostly on the publication of research articles and other scientific reports, forming the bedrock of knowledge in most fields, with rare instances of self-correction. In this prevailing culture, “[t]he thought of having to retract an article can instill fear into the heart of scientists, who see it as equivalent to being named and shamed. There are currently few incentives for honesty, and keeping quiet about an error will often seem the easiest option” (Bishop, 2018).

It is thus timely to investigate what factors underlie retractions and self-retractions for honest errors. Previous studies have shown the distribution of retractions and its reasons among journals (Fang and Casadevall, 2011; Fang et al., 2012; Gasparyan et al., 2014; Vuong et al., 2020), research fields (Grieneisen and Zhang, 2012; Ribeiro and Vasconcelos, 2018; Vuong et al., 2020), and countries (Fang et at., 2012; Amos, 2014; Fanelli et al., 2015; Ribeiro and Vasconcelos, 2018). When it comes to reasons for retractions, a considerable share is attributed to misconduct, in different fields, with smaller fractions for honest errors (Fang et at., 2012; Bozzo et al., 2017; Ribeiro and Vasconcelos, 2018; Li et al., 2018; Coudert, 2019; Christopher, 2022).

Retractions and self-retractions can reveal much about the social dimension of the scientific enterprise. That said, the understanding of factors influencing this correction process should be sought in light of a research culture, including its publication system, that does not incentivize public exposure of failures (Allison et al., 2016; Bishop, 2018; Roher et al., 2021). When it comes to such exposure through self-retractions for honest error, although there have been growing efforts toward normalizing this process (Bishop, 2018; Ribeiro et al., 2022), such cultural shift takes time. One issue is perceptions among scientists that one’s reputation may be tainted in this process (Fanelli, 2016; Hosseini et al., 2018; Bishop, 2018). In fact, there are several gaps in our understanding of factors underlying the individual self-correction of science for honest error, including possible influences of fields, countries, and gender.

Concerning the latter, given gender disparities in academia, it is worth investigating whether female scientists are more (or less) proactive than male scientists towards correcting the research record for honest errors, across fields and countries. As well documented, gender disparities are part of the history of science, and they have posed several barriers that female scientists have to overcome for being recognized in academia.

An American perspective on this matter was brought by Margaret Rossiter, a well-known science historian who coined the phrase “*Matilda Effect*” (Rossiter, 1993). Different from the Matthew Effect, a biased recognition towards those who are already eminent in science (Merton, 1968), the Matilda Effect is the result of prejudice that women face in academia, leading their work to be overlooked or even credited to male colleagues (Rossiter, 1993; Lincoln et al., 2012). Female researchers themselves may have implicit gender biases against their own peers (Lincoln et al., 2012; Knobloch-Westerwick et al., 2013; Raymond, 2013).

That said, gender inequalities remain a challenge for women in science. For example, a comprehensive study on gender disparities in science across 83 countries and 13 disciplines shows that the gender gap in terms of research productivity is a widespread phenomenon (Huang et al., 2020). Female scientists secure fewer first and last authorship positions (Larivière et al., 2013; West et al., 2013; Hart and Perlis, 2019; Ross et al., 2022), compound fewer peer-review and editorial boards (Helmer et al., 2017), tend to publish in lower impact journals in some fields (Larivière et al., 2013; Molwitz et al., 2021, Bendels et al., 2018), usually receive fewer citations (Larivière et al., 2013; Bendels et al., 2018; Shamsi et al., 2022), less funding (Ley and Hamilton, 2008; Oliveira et al., 2019), fewer awards (Lincoln et al., 2013; Meho, 2021), and patents (Ross et al., 2022).

It is against this backdrop that we have explored the role of gender in self-correcting the research record. For example, this unfavorable environment for women in academia may make female authors more reluctant to self-retract research articles, even for honest errors, as they might fear the outcome. Looking at gender and retractions, Decullier and Maisonneuve (2021) found that among 120 retractions analyzed, 37.2% were authored by female authors, with male authors accounting for 59.2% for fraud and plagiarism. However, this analysis was not based on sole corresponding authors, who are expected to have a decisive role in initiating a retraction. We explored the representation of gender in self-correcting science through research articles with sole corresponding authors, based on a dataset of 3,822 retractions attributed to error.

## Methodology

We collected data on retractions classified under the category *error* from the Retraction Watch Database (01/01/2010-31/12/2021 - 12 years in total). A total of 3,822 records were obtained, with information on authorship, article type, DOI of the original publication, DOI of the retraction notice, and nature of the publication: research article, letter, case report, review article, clinical study, conference abstract, meta-analysis, preprint, commentary/editorial, book chapter, auto/biography, trade magazine, correction/erratum, guideline, governmental publication, interview, supplementary material.

We selected only research articles (RA), considering the impact of the correction of original data on the research record. After excluding any record that mentioned “investigation by” as such categorization may involve other reasons rather than honest error, we screened the dataset for retraction for error in analyses and/or data and/or methods and/or materials and/or conclusions and/or image and/or text. We excluded records that combined this information with at least one of the following: false/forged authorship, paper mill, ethical violations, and/or fake peer review, misconduct, falsification, fabrication, plagiarism, publisher or third party, concerns/issues about data or original data not provided (when retraction notice was not clear), concerns/issues about data and/or authorship and/or referencing/attributions, duplication, manipulation. This refinement was necessary to prevent that error and misconduct, or error and other unknown or even obscure reasons would be categorized as honest error. RAs with unidentified or with more than one corresponding author were excluded. We obtained 575 notices and excluded those with insufficient information in our search (n=11), with unclear or obscure reasons, with an indication that the retraction was not initiated by the authors, and with more than one corresponding author (“raw dataset”). We obtained 472 self-retractions with only one corresponding author, initially attributed to honest error. This final dataset (n= 472) included the complete names (at least first name and surname) of the corresponding author, with additional information from the Web of Science. We used the Genderize database for gender prediction, which is based on total counts of a first name and on the probability of prediction. Further details can be found at https://genderize.io/. The gender of these corresponding authors was predicted for 413 (88%) of the 472 corresponding authors included in our refined dataset. After a preliminary analysis of the content of each notice in the “raw dataset” (n=564), a total of 464 self-retractions for honest error were validated by two members of the team. From the 413 with gender predicted, a total of 350 was obtained after further refinement, but notices with gender prediction below 90% were excluded (n=69). A total of 281 valid self-retraction notices attributed to honest error were obtained and classified according to gender. **Figure 1** shows the steps of our screening scheme:

**Figure 1.**
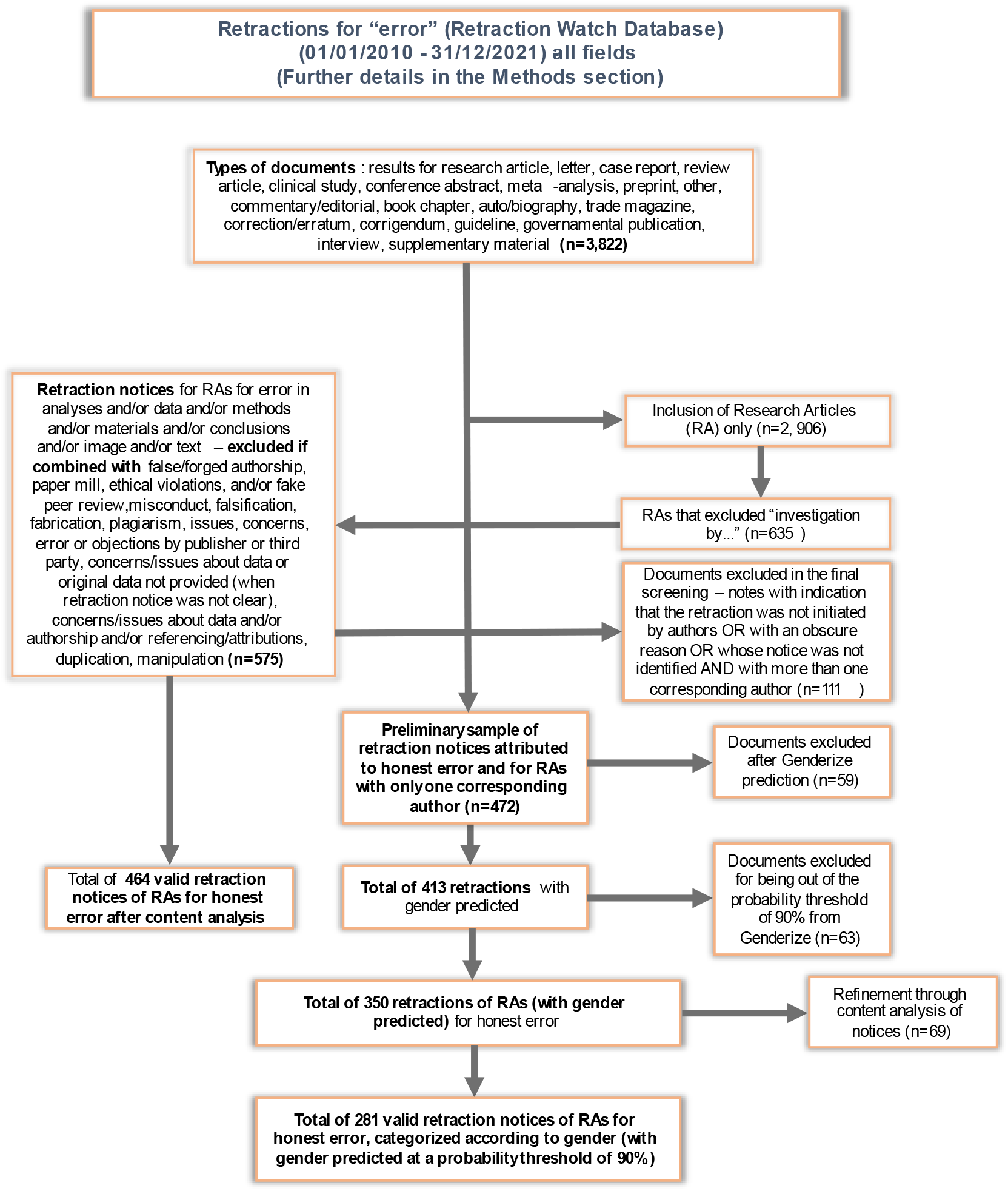
Screening scheme for the final dataset on self-retractions for honest error (n=413) of research articles with sole corresponding authors, originally extracted from 3,822 records for retractions for error (2010-2021) in the Retraction Watch Database.

## Results and Discussion

We selected all retraction notices, classified under the category *error*, of research articles from the period 2010 and 2021, collected from the Retraction Watch Database, and set up a dataset with 3,822 retraction records for error. As shown in the screening scheme in **Figure 1**, we obtained 281 self-retractions for honest error of research articles with only one corresponding author, with gender predicted with a 90% probability threshold. This number is equivalent to 61% of the total self-retractions of research articles with notices exclusively attributed to honest error (n=464) in our dataset, including those research articles that had more than one corresponding author. This number (n=464) is equivalent to 16% of the 2,906 retractions of research articles classified under the category of “error” in the Retraction Watch Database for the period 2010-2021.

**Figure 2a** shows the distribution of valid self-retractions for honest error of research articles (n=281, with only one corresponding author) across fields, according to Journal Citation Reports (JCR), with gender predicted by Genderize, with 25% female- and 75% male-authored records. **Figure 2b** offers an overview of the distribution of probability for the name of the corresponding being female or male.

**Figure 2a.**
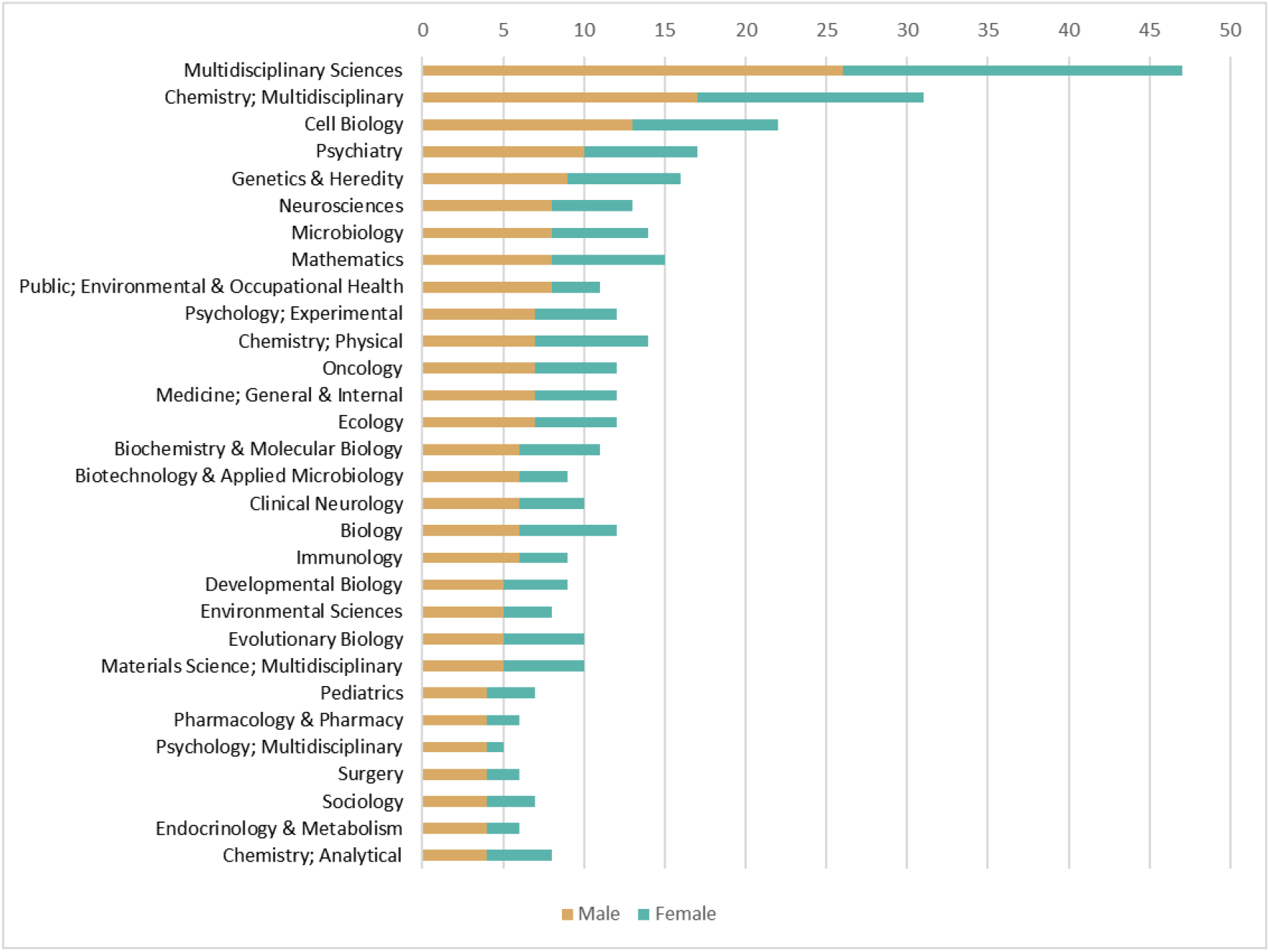
Distribution across fields of corresponding authors in terms of gender (female X male) of valid self-retractions for honest error of research articles (n=281) recorded in the Retraction Watch Database (2010-2021), with female corresponding authors accounting for 25% of these records. The data are plotted for the 30 most frequent categories.

**Figure 2b.**
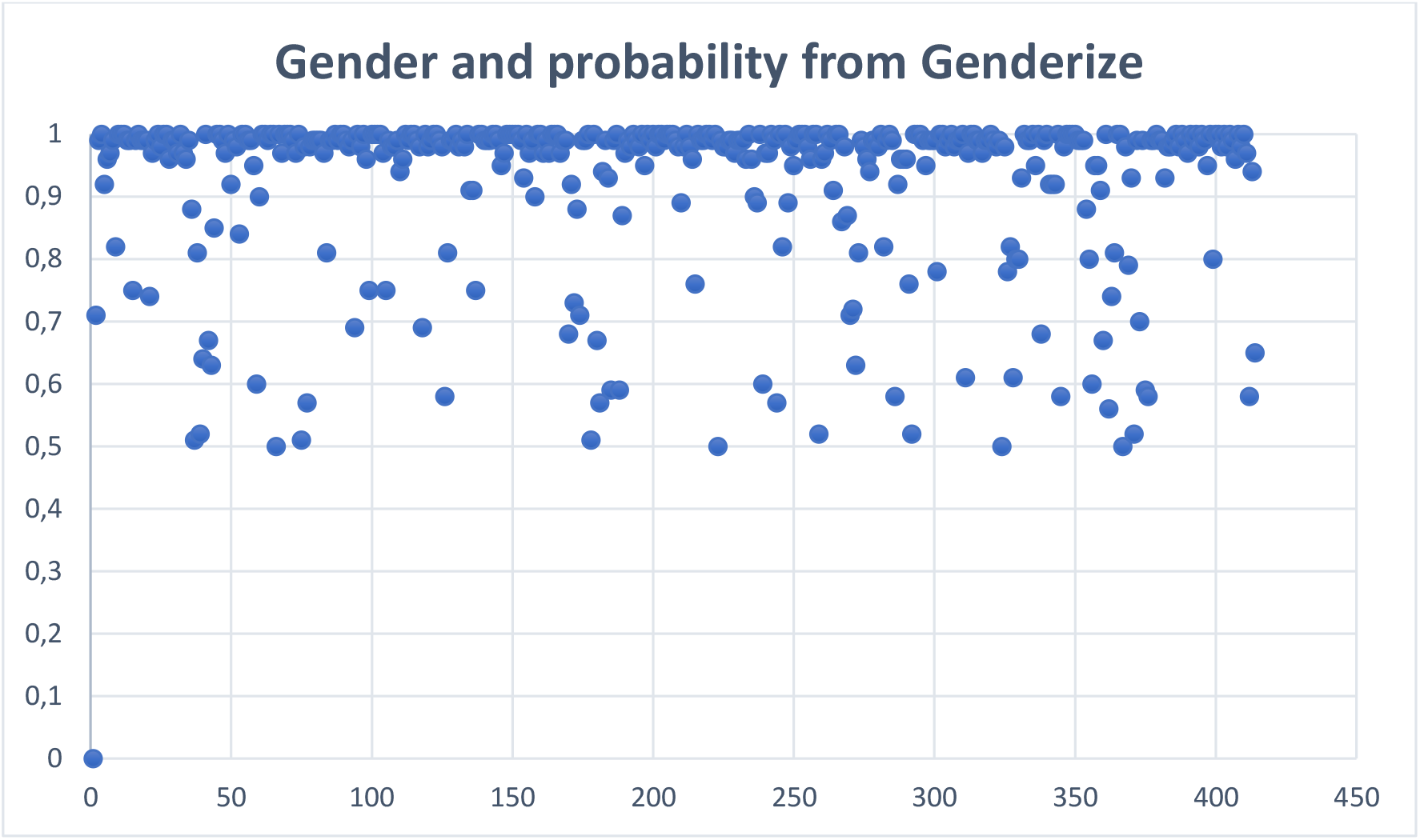
Distribution of prediction probability (between 50% and 100%) for female and male names of corresponding authors of research articles self-retracted (n=413), predicted by Genderize.

The results show that 25% (n=71) of valid self-retraction notices of research articles for error (n=281) (2010-2021) were led by female scientists, who were the sole corresponding authors of the research articles. A previous study exploring characteristics, global distributions, and reasons for 1,339 retractions from PubMed and Retraction Watch website showed that “[f]or all reasons of retraction, the percentage of retracted articles with male senior or corresponding authors was substantially higher than that with female senior or corresponding authors” (Li et al., 2018, p. 41). As for retractions for error, the same authors reported that female corresponding authors accounted for 19.2% (n=37) of the total of retractions attributed to error (n=193) (Li et al., 2018). Decullier and Maisonneuve (2021) investigated the underrepresentation of women in retractions and identified the reasons for 113 retractions for female and male authors and found that 37.2% retractions were for publications first authored by female scientists. The study also showed that reasons for retraction differed considerably comparing female and male authors, with 28.6% of retractions for research misconduct for female and 59.2% for male authors (Decullier and Maisonneuve, 2021). These percentages are consistent with evidence brought by Fang et al. (2013), who revealed that male scientists were overrepresented (about two thirds of 228 individuals) among those committing research misconduct in the life sciences.

Drawing upon Pohlhaus et al. (2011) and Nosek et al. (2007), Kaatz et al. (2013) reported that “[i]f we use NIH research award dollars as a proxy for the opportunity to commit fraud in the life sciences, we find that men have substantially more opportunity to commit fraud than women. Compared to women, men are more likely to hold multiple simultaneous R01 awards, lead large center grants, and successfully compete when submitting renewals (20–22)” (Kaatz et al., 2013, p.2).

This overrepresentation of male scientists is also marked in our dataset, across fields (**Figure 3**). In this figure, each corresponding author is associated to the article field (gray vertex). The size of each vertex is proportional to the number of authors associated with it. In order to simplify the data visualization, **Figure 3** displays only the two largest connected components of the generated network.

**Figure 3.**
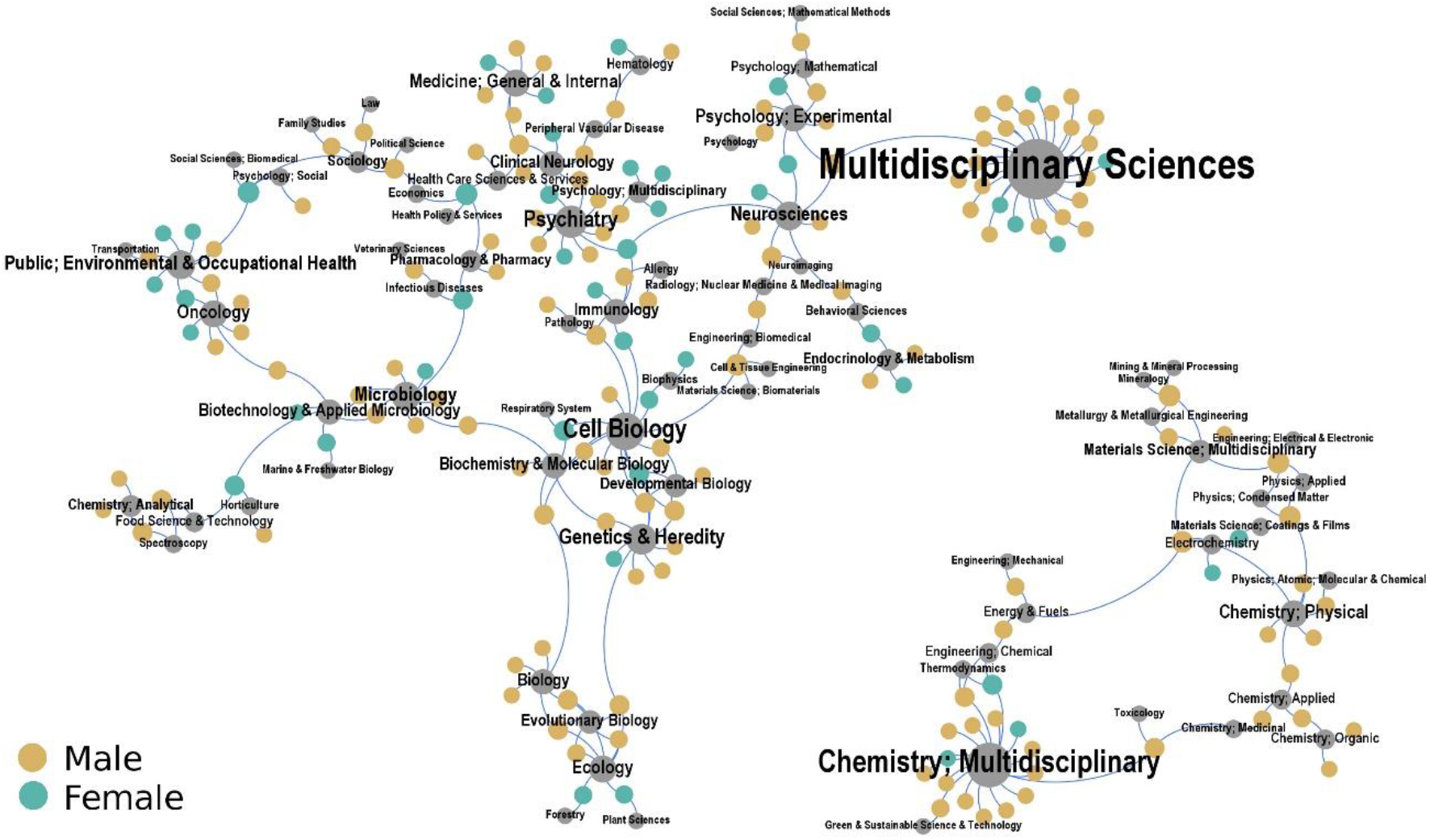
Network of self-retractions for honest error, considering research articles (n=244), across fields (according to JCR), recorded in the Retraction Watch Database (2010-2021), for sole corresponding authors, female or male. Note that the fields for 37 documents in our final dataset were not identified.

Figure 3 shows self-retractions for honest error by male corresponding authors, compared to female authors, distributed across the fields. As can be seen, male scientists are prevalent for most of these fields, including multidisciplinary and chemistry multidisciplinary (83%, with 34 out 41 records for these fields), and the medical, biomedical, life (including environmental) and health sciences (67%, with 96 out 143 records for these fields). The prevalence of self-retractions of male corresponding authors in these latter fields might suggest that they are more proactive than female corresponding authors towards this type of correction. Nevertheless, the high-impact factor of most journals (**Table 1**) may be a confounder. Note, for example, that male corresponding authors account for 82% (n=46) of 56 research articles with valid self-retractions for honest error, published in journals with impact factors between 10 and 176.

High-impact factor journals would tend to correct more (Fang and Casadevall, 2011; Fang et al., 2012; Brainard, 2018) than the other journals. Additionally, the fact that the medical, biomedical, life, and health sciences have taken the lead in the discussion of publication ethics is likely to influence this pattern.

**Table 1.** List of 46 research articles (82%), out a total of 56, with valid self-retractions for honest error authored by male scientists, published in journals with impact factors between 10 and 176 (n=56), according to JCR.

| Fields - according to Journal Citation Reports (JCR) | Impact Factor (IF) |
| --- | --- |
| Multidisciplinary Sciences - Scie(Q1) | 12.779 |
| Oncology - Scie(Q1) | 11.816 |
| Medicine, General & Internal - Scie(Q1) | 176.079 |
| Genetics & Heredity - Scie(Q1); Developmental Biology - Scie(Q1); Cell Biology - Scie(Q1) | 12.89 |
| Cell Biology - Scie(Q1); Biochemistry & Molecular Biology - Scie(Q1); Biology - Scie(Q1) | 10.9 |
| Rheumatology - Scie(Q1) | 27.973 |
| Ophthalmology - Scie(Q1) | 14.277 |
| Psychiatry - Scie(Q1); Psychiatry - Ssci(Q1) | 25.911 |
| Medicine, General & Internal - Scie(Q1) | 176.079 |
| Psychology, Multidisciplinary - Ssci(Q1) | 10.172 |
| Multidisciplinary Sciences - Scie(Q1) | 12.779 |
| Chemistry, Multidisciplinary - Scie(Q1); Green & Sustainable Science & Technology - Scie(Q1) | 11.034 |
| Multidisciplinary Sciences - Scie(Q1) | 63.714 |
| Immunology - Scie(Q1); Allergy - Scie(Q1) | 14.29 |
| Immunology - Scie(Q1); Allergy - Scie(Q1) | 14.29 |
| Chemistry, Multidisciplinary - Scie(Q1) | 16.383 |
| Neurosciences - Scie(Q1); Immunology - Scie(Q1); Psychiatry - Scie(Q1) | 19.227 |
| Chemistry, Multidisciplinary - Scie(Q1) | 24.274 |
| Peripheral Vascular Disease - Scie(Q1); Clinical Neurology - Scie(Q1) | 10.17 |
| Clinical Neurology - Scie(Q1) | 11.8 |
| Multidisciplinary Sciences - Scie(Q1) | 63.714 |
| Psychiatry - Scie(Q1); Psychiatry - Ssci(Q1) | 25.911 |
| Psychiatry - Ssci(Q1); Psychiatry - Scie(Q1) | 11.225 |
| Ecology - Scie(Q1); Evolutionary Biology - Scie(Q1) | 19.1 |
| Oncology - Scie(Q1) | 33.006 |
| Multidisciplinary Sciences - Scie(Q1) | 12.779 |
| Chemistry, Multidisciplinary - Scie(Q1) | 16.823 |
| Multidisciplinary Sciences - Scie(Q1) | 63.714 |
| Multidisciplinary Sciences - Scie(Q1); Psychology, Experimental - Ssci(Q1); Neurosciences - Scie(Q1) | 24.252 |
| Multidisciplinary Sciences - Scie(Q1) | 12.779 |
| Multidisciplinary Sciences - Scie(Q1) | 12.779 |
| Multidisciplinary Sciences - Scie(Q1) | 12.779 |
| Medicine, General & Internal - Scie(Q1) | 157.335 |
| Chemistry, Multidisciplinary - Scie(Q1) | 16.383 |
| Cell Biology - Scie(Q1); Developmental Biology - Scie(Q1) | 13.417 |
| Multidisciplinary Sciences - Scie(Q1) | 63.714 |
| Chemistry, Physical - Scie(Q1) | 13.7 |
| Environmental Sciences - Scie(Q1); Engineering, Environmental - Scie(Q1) | 11.357 |
| Biochemistry & Molecular Biology - Scie(Q1); Cell Biology - Scie(Q1) | 12.067 |
| Multidisciplinary Sciences - Scie(Q1) | 12.779 |
| Chemistry, Multidisciplinary - Scie(Q1) | 16.823 |
| Medicine, General & Internal - Scie(Q1) | 44.409 |
| Chemistry, Multidisciplinary - Scie(Q1) | 24.274 |
| Anesthesiology - Scie(Q1) | 12.893 |
| Hematology - Scie(Q1) | 25.476 |
| Hematology - Scie(Q1) | 25.476 |
| Hematology - Scie(Q1); Peripheral Vascular Disease - Scie(Q1) | 10.514 |
| Genetics & Heredity - Scie(Q1) | 11.043 |
| Genetics & Heredity - Scie(Q1); Developmental Biology - Scie(Q1); Cell Biology - Scie(Q1) | 12.89 |
| Genetics & Heredity - Scie(Q1); Developmental Biology - Scie(Q1); Cell Biology - Scie(Q1) | 12.89 |
| Environmental Sciences - Scie(Q1) | 10.753 |
| Pediatrics - Scie(Q1) | 26.796 |
| Medicine, General & Internal - Scie(Q1) | 157.335 |
| Endocrinology & Metabolism - Scie(Q1) | 44.867 |
| Multidisciplinary Sciences - Scie(Q1) | 63.714 |
| Multidisciplinary Sciences - Scie(Q1) | 63.714 |

Overall, and in accordance with previous works already cited in this section, our results indicate that self-retractions for honest error are mostly male led. We categorized the retractions for honest error in our dataset according to country and found that 87 records (31% of the total of 281) were from sole corresponding authors affiliated with institutions in the United States. The country accounts for 41% of all female scientists (n=71) in our dataset with gender predicted (n=281) (**Figure 4**).

**Figure 4.**
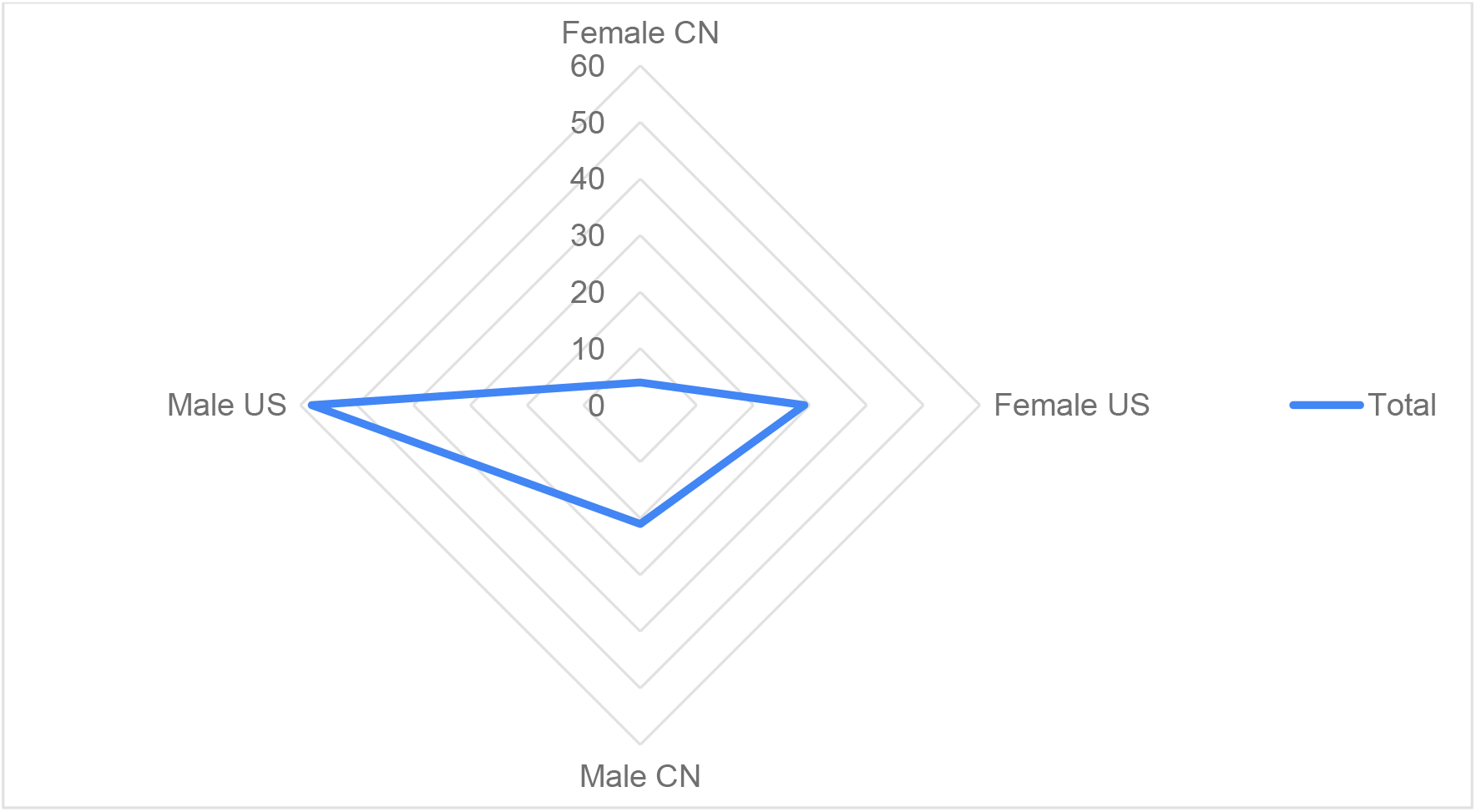
Distribution of self-retractions for honest error (n= 112) of research articles recorded in the Retraction Watch Database (2010-2021), for sole corresponding authors affiliated with institutions from the United States (n=87) and China (n=25), with 33% (n=29), and 16% (n=4) female-authored notices.

On the one hand, these data do not allow us to infer that female scientists affiliated with the two most productive countries in terms of publication output (National Science Foundation, 2021) have taken a more proactive role in self-correcting science for honest error. In addition to the small size of our sample, there might be “false positives” – for example, some of these corresponding authors who retracted the paper for honest error could have done that not by their own initiative but by a request from editors or a third party. On the other hand, and despite these caveats, these results are at least intriguing. Considering that the United States is one of the major countries leading discussions and actions toward addressing gender disparities in science in the last decades, these data might be interpreted as reflecting this factor. Concerning China, gender disparities have a long history and continue to be challenging, although great strides have been taken in the last decades (Gu, 2021).

As gender biases and disparities have been increasingly recognized as sources of damage for the career of researchers and for the research enterprise at large, in many countries and fields, these results add another layer to the growing body of literature addressing the influence of gender issues in the publication system for female researchers, across countries. In this publication arena, Roher et al. (2021, p.1265) note that “researchers may often be reluctant to initiate a retraction given that retractions occur most commonly as a result of scientific misconduct (Fang et al., 2012) and are, therefore, often associated in the public imagination with cases of deliberate fraud.” Also, individual self-corrections may imply an individual cost (Roher et al., 2021), with greater impact on the attitude of female corresponding authors facing problems in their research articles, including errors in data, results, and conclusions.

## Conclusions

Our results indicate that the percentage of self-retractions of research articles, issued in the last 12 years and attributed solely to unintended mistakes, is low (16%). Although we used rather strict criteria, this result reinforces that only a small fraction of the retracted literature can be assigned to honest error. When it comes to gender, we have found that self-retractions for honest error have been mostly male led, with prevalence for authors affiliated with institutions in the United States and China, the two most productive countries in terms of publication output. According to our results, male corresponding authors have been more proactive to self-correct the research record for unintended mistakes, accounting for 75% of the notices. One possible explanation is that these male corresponding authors have come across post-publication issues, for example, in their data, results, and/or conclusions, more often than female corresponding authors. However, these possible explanations cannot tell the whole story, considering the social dimension of retractions and the underrepresentation of female scientists in the literature as corresponding authors. The percentage is rarely higher than 30% for most fields (Bello and Galindo-Rueda, 2018; Son and Bell, 2022). A possible relationship of our results with this overall picture merits further investigation.

As we had suggested previously, the perception that retractions would taint the reputation of scientists might be stronger among women, which may be a source of unconscious bias in this correction process. After all, “[e]xisting recognition and reward structures offer no external incentive to come forward and request a retraction of your paper upon discovering a fatal honest error.” (Nature Human Behavior, 2021, p.1591). As social structures in academia are entangled with gender disparities, whether such disparities have played a role in discouraging female scientists, at different career stages, to come forward and correct the research record for honest error through self-retractions is a wide-open question.

## Limitations

Our study has several limitations worth noting. First, the source of the data is subject to research material that is not free from bias – the Retraction Watch Database records information from retraction notices whose content is not necessarily detailed and may involve overlapping classifications. For example, not everything classified under the category error is restricted to it as retractions can include error and other issues not always detailed by editors and/or authors. That said, our category “honest error”, although resulting from a careful screening and independent crosschecking of the notices, relies mostly on the honesty of the authors. Whereas we applied stringent criteria to include honest-error notices in our sample, we cannot take for granted that all these notices are overly honest and/or bias-free reports.

Second, we adopted a binary (female or male) category for gender, which is the only possible given restrictions imposed by the way the publication system is organized so far. The third issue is the threshold used for gender prediction, obtained from Genderize, which is conservative, as of 90%. Yet, a less conservative threshold – starting at 75%, for example, leads to an increment of only 4% in the representation of female corresponding authors. One additional caveat is the size of our sample of valid self-retractions for honest error with reliable gender prediction (n=281), equivalent to 61% of the 464 valid self-retraction notices for honest error obtained.

Despite this caveat, we set up strict criteria for a notice to be considered a self-retraction and exclusively attributed to error. We thus believe, on the basis of such criteria, that our results offer a reliable picture of the representation of gender (female X male) in the self-correction of science for honest error through retractions. Considering the crucial role of corresponding authors to help correct the research record, further studies exploring the representation of female scientists in this process are timely.

## Acknowledgments

We thank Professor Gabriella Andrea de Castro Pérez, at the Federal Institute of Education, Science and Technology of Rio de Janeiro (IFRJ), for her critical reading of this manuscript and relevant comments.

